# Influence of the ATP-dependent DNA ligase, Lig E, on *Neisseria gonorrhoeae* microcolony and biofilm formation

**DOI:** 10.1101/2025.02.17.638724

**Authors:** Jolyn Pan, Abdullah Albarrak, Joanna Hicks, David Williams, Adele Williamson

## Abstract

**Background:** *Neisseria gonorrhoeae*, the causative agent of the sexually transmitted infection, gonorrhoea, is known to form biofilms rich in extracellular DNA on human cervical cells. Biofilm formation is conducive to increased antimicrobial resistance and evasion of the host immune system, potentially causing asymptomatic infections. Using plate-based assays we have previously shown that disruption of a potential extracellular DNA ligase, Lig E, in *N. gonorrhoeae* (Ngo-Lig E) impacts biofilm formation. In this research, we further explored this phenotype using confocal and scanning electron microscopy to directly visualise the morphology of microcolony and biofilm formation. Biofilm growth on artificial surfaces and on 3-dimensional human vaginal epithelial tissue was evaluated for strains where *ngo-lig E* was either disrupted or overexpressed.

**Results:** Results demonstrated that Ngo-Lig E was important for the formation of robust, compact *N. gonorrhoeae* microcolonies, as well as extensive biofilms on artificial surfaces. The *ngo-lig E* deletion strain also had the highest tendency to be retained on the surface of epithelial tissues, with decreased invasion and damage to host cell layers.

**Conclusions:** These findings support a role for Ngo-Lig E to be secreted from *N. gonorrhoeae* cells for the purpose of inter-cell adhesion and biofilm formation. We suggest that Ngo-Lig E strengthens the extracellular matrix and hence microcolony and biofilm formation of *N. gonorrhoeae* by ligation of extracellular DNA.

## Introduction

*Neisseria gonorrhoeae* is a Gram-negative diplococcus bacterium responsible for the sexually transmitted infection (STI), gonorrhoea. With the World Health Organisation estimating 106 million new cases each year, gonorrhoea is the second most common STI in the world (Unemo & Shafer, 2014; World Health Organization, 2024). Infections occur in the mucosal epithelial cells of the urogenital tract, causing inflammation that presents as pain with urination in men (urethritis) and abnormal bleeding and pain in women (cervicitis) (Edwards & Apicella, 2004; Tapsall, 2005). Additionally, infections in women may spread to the upper reproductive tract, causing pelvic inflammatory disease, as well as ectopic pregnancy or infertility (McCormack, 1981). If left untreated, infections in pregnant women may also lead to neonatal conjunctivitis and blindness in the newborn (McCormack, 1981; Dillard, 2011). In females, a high proportion of *N. gonorrhoeae* infections are asymptomatic (≥50%), allowing the bacterium to spread undetected in the community (McCormack, 1981; Unemo & Shafer, 2014). This trait is often attributed to the bacterium’s ability to readily form biofilms, which aid in oxidative stress survival, attachment to surfaces and evasion from the host immune system (Greiner *et al*., 2005; Falsetta *et al*., 2009).

Interestingly, *N. gonorrhoeae* lacks the genes necessary to produce the exopolysaccharides that contribute to biofilm architecture in other bacterial species (Greiner *et al*., 2005). Instead, *N. gonorrhoeae* utilises extracellular DNA (exDNA) as the major component which provides structural integrity to the biofilm (Steichen *et al*., 2011). This exDNA may originate from the frequent autolysis that occurs in *N. gonorrhoeae* cells, or from active secretion of DNA via the type IV secretion system (T4SS) (Morse & Bartenstein, 1974; Hebeler & Young, 1975; Elmros *et al*., 1976; Zola *et al*., 2010; Zweig *et al*., 2014). In addition, the membranous extensions or blebs of the outer membrane that are extruded during gonococcal biofilm formation often harbour DNA (Dorward *et al*., 1989; Greiner *et al*., 2005; Steichen *et al*., 2008). The abundance of exDNA in the biofilm has the potential to act as a pool for gene exchange and acquisition of new antibiotic resistance genes (Kouzel *et al*., 2015); however the extent of DNA diffusion through established gonococcal biofilm is modulated by its maturity and density, which may limit horizontal gene transfer by this mechanism (Kouzel *et al*., 2015; Bender *et al*., 2022).

The DNA component of gonococcal biofilms is enzymatically remodelled by a secreted thermonuclease, Nuc, that degrades exDNA in the biofilm matrix, as well neutrophil extracellular traps (NETs), the latter aiding in bacterial escape from NET killing (Steichen *et al*., 2011; Juneau *et al*., 2015). However in addition to this nuclease, *N. gonorrhoeae* also encodes a minimal ATP-dependent DNA ligase, Lig E, which like Nuc, possesses an N-terminal signal peptide that is predicted to direct its extracellular secretion. Removal of this signal sequence has been shown to increase both stability and activity of recombinantly-expressed Lig E, which promotes the view that it is the cleaved isoform that represents the biologically-relevant mature protein. Lig E is encoded in the genomes of many Gram-negative bacteria without any synthetic organisation or consistent co-localisation with other genes (Williamson *et al*., 2016; Pan *et al*., 2021). This, together with the presence of the N-terminal signal sequence, suggests a function of Lig E other than chromosomal DNA repair and that it may act on exDNA (Magnet & Blanchard, 2004; Williamson *et al*., 2014). Consistent with this is the fact that Lig E is found in many biofilm-forming and competent proteobacteria like *N. gonorrhoeae* (Williamson *et al*., 2016). Recently, we reported that deletion of *lig E* from *N. gonorrhoeae* (*ngo-lig E*) negatively impacted the extent of biofilm formation when measured via an indirect crystal violet assay, as well as impacting *N. gonorrhoeae* adhesion to host human cervical cells (Pan *et al*., 2024).

In the present study, we further explore this phenotype, visualising the biofilms formed by *ngo-lig E* deletion and overexpressing strains of *N. gonorrhoeae* using microscopy. We provide the first reported use of the Centre for Disease Control (CDC) Biofilm Reactor® (BioSurface Technologies) to generate constant shear forces during growth of *N. gonorrhoeae* biofilms, in conjunction with confocal laser scanning microscopy (CLSM). We also used commercially available 3-dimensional (3-D) reconstructed human vaginal epithelium (rHVE) (SkinEthic Laboratories) in lieu of traditional 2-dimensional (2-D) cell lines for host-cell assays. Such reconstructed epithelial models account for different cell morphologies, tissue architecture and differentiation that occur during normal microbial infection *in* vivo (Malfa *et al*., 2023). Use of these approaches demonstrated the potential importance of Ngo-Lig E on *N. gonorrhoeae* microcolony and biofilm formation, as well as quantifying the damage to human tissue.

## Methods

### *Neisseria gonorrhoeae* manipulation

All *N. gonorrhoeae* used in this study were of the MS11 strain (GenBank: CP003909.1). Gonococci were grown at 37°C with 5% CO_2_ either on gonococcal base (GCB) agar (Difco) or in gonococcal base liquid (GCBL) (15 g/L Bacto™ Protease Peptone No. 3, 4 g/L K_2_HPO_4_, 1g/L KH_2_PO_4_, 1 g/L NaCl), both supplemented with 1% *Kellogg’s* supplement (22.22 mM glucose, 0.68 mM glutamine, 0.45 mM cocarboxylase, 1.23 mM Fe(NO_3_)_3_) (Dillard, 2011). Liquid growth was supplemented with sodium bicarbonate (0.042%), while solid growth was maintained in a 5% CO_2_ atmosphere. Piliation status was determined by morphology under a dissecting microscope at the start of each experiment.

The *Δnuc*^*kan*^ mutant (Table 1) was generated in the same manner as the previously-described mutants used in this study via spot transformation (Dillard, 2011; Pan *et al*., 2024). Briefly, the transforming DNA constructs were ordered as gene fragments from Twist Biosciences with flanking sequences surrounding the *nuc* site to facilitate homologous recombination. Piliated, Opa negative (Opa-) colonies were streaked through 10 ng spots of the DNA construct. Mutants were selected on GCB agar with 50 µg/mL kanamycin before verification by PCR and sequencing.

**Table 1.** List of N. gonorrhoeae MS11 mutants used in this study (GenBank: CP003909.1)

| Mutant | Insertion | Purpose | Source |
| --- | --- | --- | --- |
| $\Delta ngo-lig E^{kan}$ | Kanamycin resistance cassette disrupting the *ngo-lig E* gene (NGFG_RS11310) | *ngo-lig E* knock-out mutant | (Pan *et al.*, 2024) |
| *ngo-lig E-his<sup>kan</sup>* | 6-His-tag and a kanamycin resistance cassette | Control for the insertion of the kanamycin resistance cassette in | (Pan *et al.*, 2024) |

| | inserted at the C-terminus<br>of *ngo-lig E* | the Δ*ngo-Lig E*<sup>kan</sup><br>mutant, with the His-tag<br>serving as an epitope tag | |
| --- | --- | --- | --- |
| *opaB-ngo-lig E*-<br>*his*<sup>kan</sup> | Codon-optimised *ngo-lig E*<br>gene under the constitutive<br>*opaB* promoter inserted in<br>a neutral site in the genome<br>(NGFG_RS15145,<br>annotated as a phage<br>protein) | *ngo-Lig E* under a strong<br>constitutive promoter in<br>a neutral site in the<br>genome to observe the<br>effects of<br>overexpression of Ngo-Lig E | (Pan *et al.*, 2024) |
| Δ*nuc*<sup>kan</sup> | Kanamycin resistance<br>cassette interrupting the<br>*nuc* (thermonuclease)<br>gene, NGFG_RS05400 as<br>outlined by (Steichen *et al.*,<br>2011) | Positive control for<br>increased biofilm<br>formation in *N.*<br>*gonorrhoeae* for<br>scanning electron<br>microscopy | This work, based<br>on (Steichen *et al.*, 2011) |

The generation of other strains used in this study (Table 1) has been reported previously (Pan *et al*., 2024). As described, the *Δngo-lig E*^*kan*^ and *wt* genomes were previously re-sequenced to confirm that there were no significant differences between the two genomes apart from the disruption of *ngo-lig E* (Pan *et al*., 2024).

### Biofilm formation in CDC Biofilm Reactors®

To facilitate imaging via confocal microscopy, *wt N. gonorrhoeae* and the *ngo-lig E* mutants (Table 1) were each transformed with the pEG2 cryptic plasmid (Christodoulides *et al*., 2000), which was kindly gifted to us by the Radcliff laboratory (University of Auckland). The pEG2 plasmid contains an *sfGFP* gene under a *porA* promoter and was introduced to the MS11 variants via spot transformation with selection via erythromycin (10 µg/mL). Successful transformants were verified by fluorescence of the sfGFP under blue light, and the plasmids were continuously maintained in *N. gonorrhoeae* by addition of erythromycin to culture media.

The pEG2 transformants of the *N. gonorrhoeae* variants were streaked and cultured for 24 h on GCB plates with erythromycin. Piliated bacteria were then lawned and cultured for 16 h on chocolate agar before resuspension in GCBL. For each mutant, a 1 mL suspension of an OD_600_ of 0.05 was used to inoculate media for growth in the CBR 90 Standard CDC Biofilm Reactor® (Biosurface Technologies), which was assembled as per the manufacturer’s protocol (Figure 1). Briefly, polycarbonate coupons (diameter: 12.7 mm; thickness: 3.8 mm) were fitted into vertical polypropylene rods in the biofilm reactor. The 1 mL inoculum was introduced via the inlet port into the vessel containing GCBL with erythromycin and sodium bicarbonate (333 mL). Batch growth with stirring was performed for 6 h before continuous flow with sterile GCBL at 0.6 mL per min for 16-17 h. The coupons were then extracted from the rods and rinsed twice in water before CLSM.

**Figure 1.**
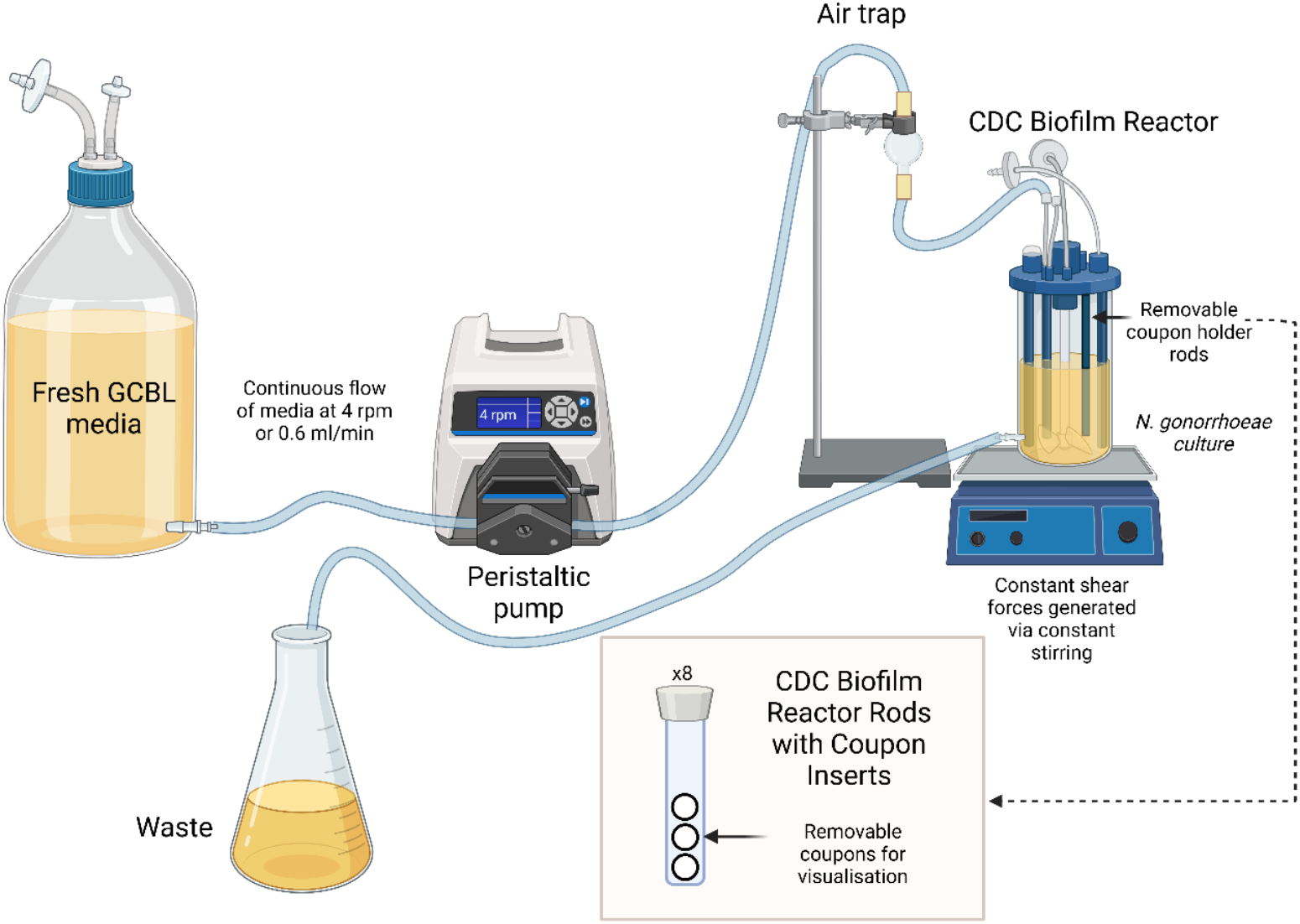
Schematic of the set up used to generate N. gonorrhoeae biofilms using the CBR 90 Standard CDC Biofilm Reactor® (Biosurface Technologies). Created in https://BioRender.com.

### Infection of reconstructed human vaginal epithelium (rHVE)

SkinEthic™ HVE tissue (0.5 cm^2^, age day 5; HVE/S/5) were ordered from Episkin (Lyon, France). These are vulval epidermoid carcinoma A431 cells seeded on a polycarbonate filter in inserts and maintained at the air-liquid interface (Figure 2). Upon receipt, cells were equilibrated with the SkinEthic™ Maintenance Medium (Episkin (Lyon, France)) for 4 h in 12-well plates in a humidified chamber (37°C, 5% CO_2_), before replacement with fresh media (1 mL). *N. gonorrhoeae* pEG2 cultures prepared in the same medium (100 µL) were inoculated onto the cells at 1×10^8^ CFU per cm^2^ and left for 16-17 h in a humidified chamber (37°C, 5% CO_2_). The supernatant under each tissue insert was recovered and used for lactase dehydrogenase (LDH) activity assays, while the cell inserts were washed twice with phosphate-bujered saline (PBS). The tissue and their membranes were then isolated from the inserts for microscopic imaging.

**Figure 2.**
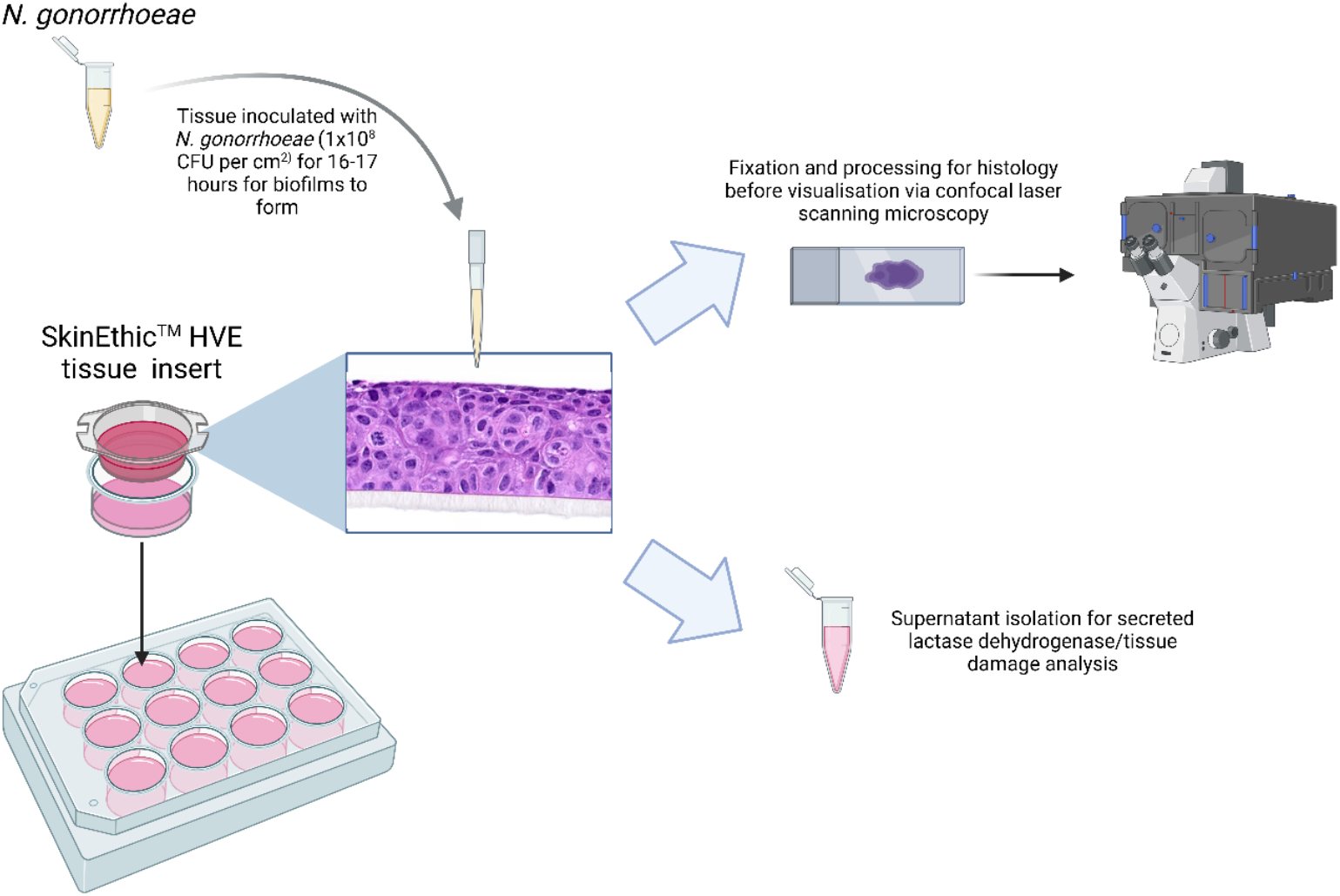
Schematic of the infection work-flow with the SkinEthic™ HVE tissue model obtained from Episkin (Lyon, France). Created in https://BioRender.com. Image of HVE cells obtained from https://www.episkin.com/HVE-Vaginal-Epithelium.

### Histological techniques

The isolated cells and membranes were individually wrapped in Surgipath® Bio-Wraps™ (Leica Biosystems) and placed in cassettes before soaking in Reagecon Formal Buffered Saline for 3 h. Dehydration was performed using the LeicaASP300S Fully Enclosed Tissue Processor (90% v/v ethanol 1 h, 95% v/v ethanol 1 h, 100% v/v ethanol 4×1 h, xylene 3×1 h). The cassettes were embedded in Surgipath® Formula ‘R’ paraffin wax (Leica Biosystems) using the Leica EG1150 Modular Tissue Embedding Center. Transverse sectioning was performed at the BioImaging Hub at the School of Biosciences at Cardiff University to obtain 20 µm sections on microscope slides. Wax was removed with xylene (5 min) before washes with 70% v/v ethanol (5 min) and 100% v/v ethanol (5 min), followed by rehydration in water (5 min). VECTASHIELD® Antifade Mounting Medium with DAPI (H-1200-10, Vector Laboratories) (20 µL) was added onto the sections before visualisation using CLSM.

### Confocal laser scanning microscopy (CLSM)

CLSM was conducted at the Cardiff University Bioimaging Hub Core Facility (RRID:SCR_022556). Biofilms grown on polycarbonate coupons were visualised using the Zeiss Cell Discoverer 7 microscope at x40 magnification using the suGFP channel (laser excitation (exc) wavelength: 480 nm, laser emission wavelength (em): 505 nm) to image the sfGFP-expressing gonococcal cells. Infected rHVE tissues were visualised using the Zeiss LSM 880 with Airyscan microscope at x63 magnification. rHVE nuclei were visualised using the DAPI channel (exc: 405 nm, em: 449 nm) and sfGFP-expressing *N. gonorrhoeae* were visualised using the GFP channel (exc: 488 nm, em: 519 nm). Five random fields-of-view (z-stacks) were obtained for each slide/coupon.

Images were analysed using COMSTAT 2.1 (Heydorn *et al*., 2000; Vorregaard, 2008) in an OME-TIFF format (Otsu thresholding). Quantified parameters included biovolume/biomass (volume over area, µm^3^/ µm^2^), average thickness (biomass) (height distribution of biomass-containing columns, µm), average thickness (entire area) (height distribution of the biofilm for the entire observed area including empty columns, µm), maximum thickness (highest point of the biofilm ignoring empty voxels, µm), surface to biovolume ratio of the biofilms (total surface facing the void over biovolume, µm^2^/ µm^3^) and dimensionless roughness coefficient, 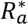 (variability in biofilm height; 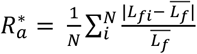 where *N*=number of measurements, *L*_*fi*_ = *i*’th individual thickness measurement and 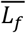 =average thickness) (Nowicki, 1985; Murga *et al*., 1995; Heydorn *et al*., 2000; Vorregaard, 2008).

To ensure consistency in acquisition of parameters, all confocal images using the Zeiss LSM 880 with Airyscan microscope were acquired at the same zoom setting (zoom setting 3), which in the infection experiments, focused on the biofilm layer at the surface of the rHVE tissue. As described below, differences in tissue thickness were observed after inoculation with didderent strains, meaning that the lower membrane was not visible in some fields-of-view. To account for this difference, the depth of *N. gonorrhoeae* invasion was normalised relative to the depth of remaining tissue in the same field-of view.

### Scanning electron microscopy (SEM)

*N. gonorrhoeae* variants were lawned on GCB agar for 16 h before resuspension in GCBL. The bacteria were seeded into 12-well plates with a starting OD_600_ of 0.05 and on to 0.2 µM pore size filter papers. After 9 h (exponential phase, based on growth experiment from Pan *et al*., 2024) the medium was removed and the bacteria fixed overnight in 2.5% glutaraldehyde. The bacteria were washed (x4) in 0.1 M sodium cacodylate and distilled water before successive dehydration with 50% ethanol (1 h), 75 % ethanol (1 h), 95% ethanol and four rinses with 100% ethanol. The critical point drying process and coating with platinum (5 nm) was performed by the Electron Microscope Facility at the University of Waikato. Images were obtained via the Hitachi SU8230 microscope (3 kV acceleration). Three fields-of-view were taken for each sample. The area of microcolonies formed (µm^2^) was quantified via ImageJ (Schneider *et al*., 2012).

### Lactase dehydrogenase (LDH) activity assay

LDH quantification was performed on the isolated rHVE supernatant after *N. gonorrhoeae* infection using the CyQUANT™ LDH Cytotoxicity Assay kit as per the manufacturer’s instructions. Absorbances at 490 and 680 nm were measured and the background absorbance at 680 nm was subtracted from that at 490 nm.

### Statistical methods

Statistical analyses of were preformed using the GraphPad Prism 9.4.0 software (https://www.graphpad.com/). One-way analysis of variance (ANOVA) with Tukey’s multiple comparisons test was used to compare the different measurements and *p* values < 0.05 were deemed statistically significant.

## Results

### Adhesion and biofilm formation of *Neisseria gonorrhoeae* on polycarbonate surfaces is dependent on Ngo-Lig E

To determine if Ngo-Lig E impacted the ability of *N. gonorrhoeae* to adhere to and to form biofilms on abiotic surfaces, their growth on polycarbonate coupons after 16-17 h cultivation in CDC Biofilm Reactors® under constant shear forces was studied. Confocal images of the coupons (Figure 3 and Figure S1) indicated that *N. gonorrhoeae* had relatively low adherence to the polycarbonate surfaces compared with other organisms that we have studied using a similar set-up, such as *Candida albicans* and *Enterococcus faecalis* (Nassar *et al*., 2023).

**Figure 3.**
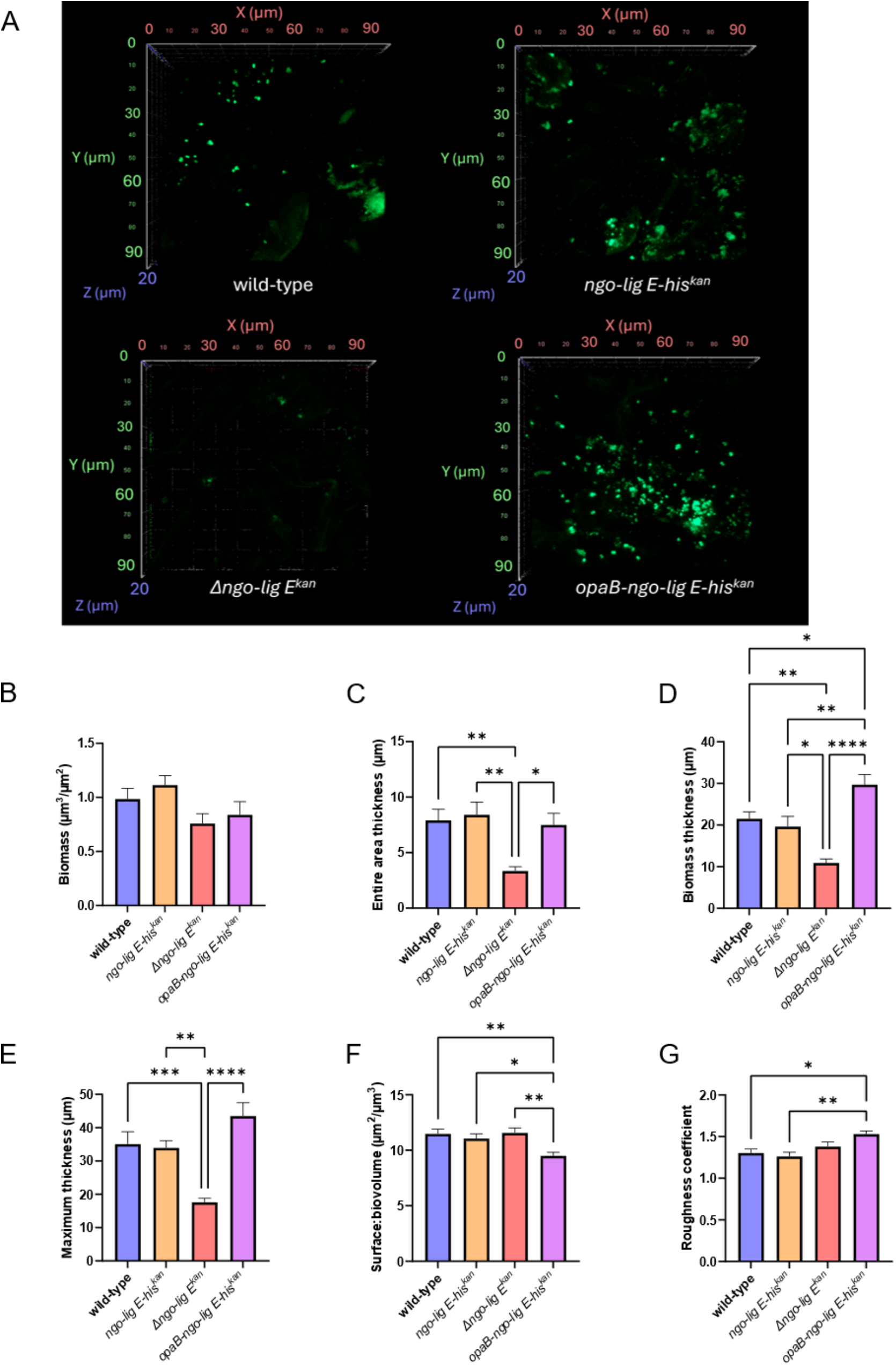
Biofilm formation and adhesion of N. gonorrhoeae pEG2 (expressing sfGFP) on polycarbonate coupons in CDC Biofilm Reactors® (a) Representative CLSM z-stack images (x40 objective magnification, em:480 nm, exc:505 nm). Five fields-of-view were imaged for each of the three biological replicates. Additional supporting images can be found in Fig S1 (b) Biomass (c) Surface to biovolume ratio (d) Dimensionless roughness coefficient (e) Average thickness (entire area) (f) Average thickness (biomass) (g) Maximum thickness. Parameters were calculated using the COMSTAT 2.1 software (Heydorn et al., 2000; Vorregaard, 2008). Points in the bar graphs are the mean of values from the five fields-of-view of three biological replicates and error bars represent the standard error of the mean. Significance values are given as * p ≤ 0.05; ** p ≤ 0.01; *** p ≤ 0.001; ** **p ≤ 0.0001. Comparisons which showed no significant differences (p > 0.05) are not indicated.

Despite this, CLSM images of the coupons after growth showed a clear decrease in the ability of the *Δngo-lig E*^*kan*^ mutant to attach to surfaces and to form extensive or continuous biofilms compared to *wt* (Figure 3 (a) and Figure S1). Conversely, biofilms formed when *ngo-lig E* was overexpressed (*opaB-ngo-lig E-his*^*kan*^) were more extensive across the surface, while *wt* and *ngo-lig E-his*^*kan*^ seemed to extend similarly to each other. Quantification via COMSTAT analysis showed no significant differences in the total biomass among the different gonococcal variants (volume per area, Figure 3 (b)). However, there was a significant reduction in both the thickness of the entire area of growth (indicative of spatial size of the biofilm across the entire area, (Figure 3 (c)) and the thickness of the biomass (thickness distribution of only biomass-containing columns, Figure 3 (d)) when *ngo-lig E* was disrupted compared to *wt* and *ngo-lig E-his*^*kan*^. Although the overexpressing *opaB-ngo-lig E-his*^*kan*^ mutant had a significantly higher average biomass thickness compared to the other three variants (Figure 3 (d)), its average thickness or spatial spread over the entire area was similar to that of *wt N. gonorrhoeae* (Figure 3 (c)), while also displaying higher overall maximum biofilm thickness (highest point of the biofilm, Figure 3 (e)) and lower surface:biovolume ratio (ratio of total surface facing the void over biovolume, Figure 3 (f)) than the other variants. Furthermore, the dimensionless roughness coefficient indicated slightly higher roughness or variability in the height of the biofilms formed by the *opaB-ngo-lig E-his*^*kan*^ mutant compared to the other *N. gonorrhoeae* variants (Figure 3 (g)).

### Ngo-Lig E increases the damage and migration of *Neisseria gonorrhoeae* into reconstructed human vaginal epithelium (rHVE) tissue

To further explore the impact of this sparse biofilm phenotype of the *N. gonorrhoeae ngo-lig E* deletion strain on pathogenicity and virulence, we investigated its ability to form biofilms on a 3-D SkinEthic™ HVE tissue model. The intended experiments involved allowing the *N. gonorrhoeae* pEG2 cells to establish biofilms on the polycarbonate coupons in CDC Biofilm Reactors® before placing these in direct contact with the rHVE tissues. However, as extensive biofilms were not formed on the coupons, we inoculated cultures of the *N. gonorrhoeae* pEG2 strains directly on to the rHVE cells to allow them to form stable biofilms on a more biologically relevant surface. Confocal imaging (Figure 4 (a) and Figure S2) of these infections showed increased depths of invasion of *wt N. gonorrhoeae*, the his-tagged mutant (*ngo-lig E-his*^*kan*^) and the overexpressor (*opaB-ngo-lig E-his*^*kan*^) in the tissue model, while *Δngo-Lig E*^*kan*^ remained on the upper surface of the tissue. Furthermore, tissues infected with the *Δngo-lig E*^*kan*^ mutant appeared more intact after inoculation, while those infected by the other *N. gonorrhoeae* variants appeared more damaged or were perforated. The final z-stack images obtained for quantification via COMSTAT were focused and zoomed onto the surface of the rHVE cells where *N. gonorrhoeae* was predicted to form biofilms (zoom setting 3). This was an optimal setting for *wt N. gonorrhoeae*, the his-tagged mutant (*ngo-lig E-his*^*kan*^) and the overexpressor (*opaB-ngo-lig E-his*^*kan*^) as it showed the extent of the damage caused by the bacteria to the host cells which decreased tissue thickness and hence the bottom membrane was visible for most. However, as a majority of the *Δngo-Lig E*^*kan*^-infected tissue were more intact, the membrane was only visible on a lower zoom setting (Figure S3).

**Figure 4.**
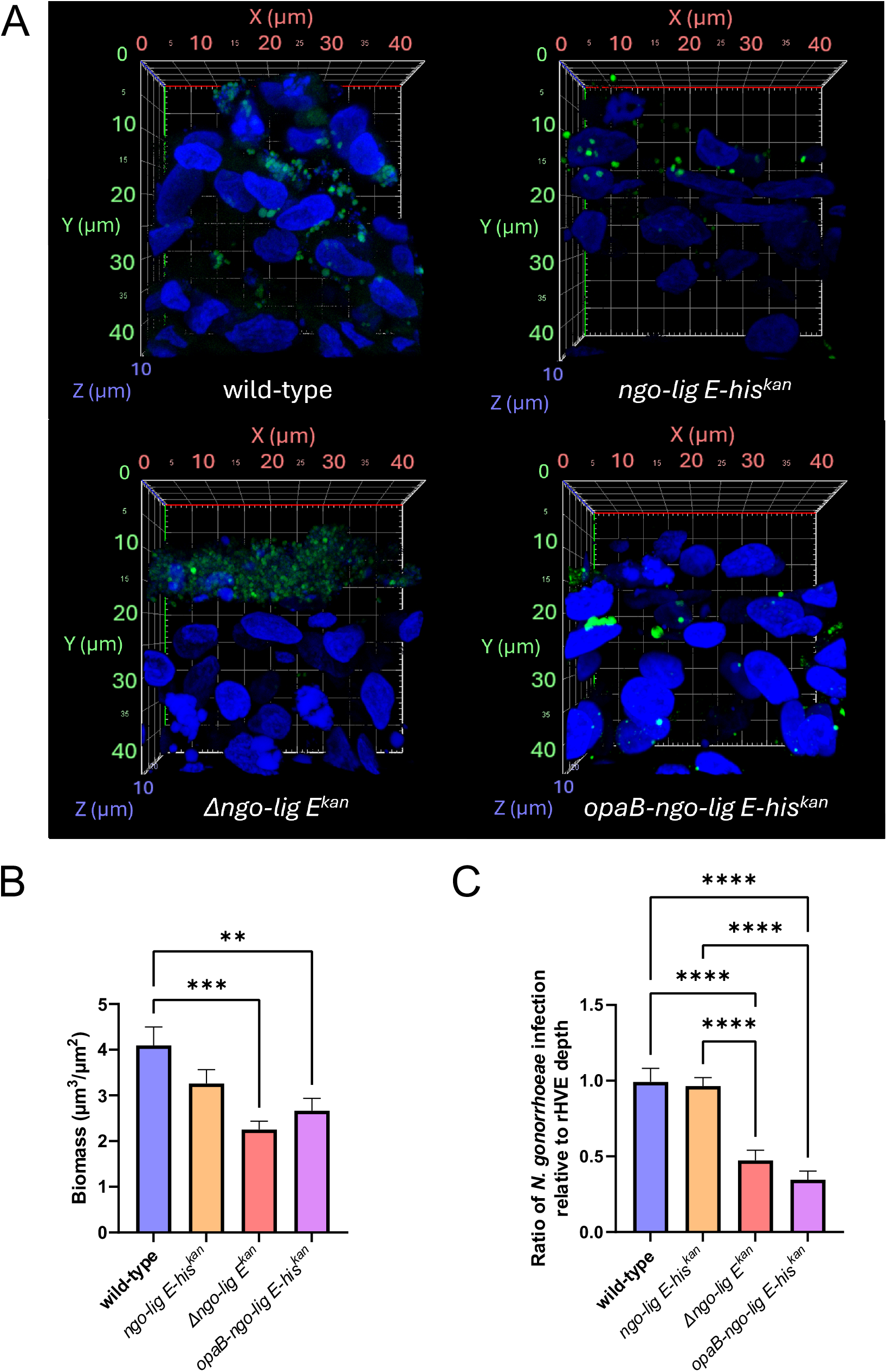
Infection and invasion of SkinEthic™ HVE cells by N. gonorrhoeae (a) CLSM z-stack images (x63 objective magnification, zoom setting 3) of the N. gonorrhoeae-infected HVE cells (20 µm) with N. gonorrhoeae pEG2 (expressing sfGFP) shown in green (exc: 488 nm, em: 519 nm) and the nuclei of the rHVE cells in blue (exc: 405 nm, em: 449 nm). Additional supporting images can be found in Fig S2 (b) Biomass of N. gonorrhoeae growth (green channel) quantified using the COMSTAT 2.1 software (Heydorn et al., 2000; Vorregaard, 2008) (d) Quantification of the depth of N. gonorrhoeae infection on the y-axis (green channel) relative to the depth of remaining HVE cells on the y-axis (blue) in the same field-of-view. Points in the bar graphs are the mean of values from the five fields-of-view of three biological replicates, with two 20 µm sections each (total 30 field-of-views). Error bars represent the standard error of the mean. Significance values are given as ** p ≤ 0.01; *** p ≤ 0.001; **** p ≤ 0.0001. Comparisons which showed no significant differences (p > 0.05) are not indicated.

COMSTAT quantification of the biofilms formed by *N. gonorrhoeae* on the rHVE cells showed a slightly lower biomass for *Δngo-Lig E*^*kan*^ and the overexpressor *opaB-ngo-lig E-his*^*kan*^ compared to *wt* on the host tissue cells (Figure 4 (b)). However, this did not take into account the extent of tissue damage induced by the bacteria. To measure this, the range of *N. gonorrhoeae* infection (highest to lowest point on the y-axis), relative to the height of remaining HVE in that particular field-of-view (Figure 4 (c)) was calculated. Results showed that both *Δngo-Lig E*^*kan*^ and *opaB-ngo-lig E-his*^*kan*^ had significantly lower invasion rates than *wt* and *ngo-lig E-his*^*kan*^, the latter two strains having similar depth ratios.

To further quantify the extent of epithelial cell damage, the amount of LDH released in the supernatant after biofilm establishment was measured as a proxy for epithelial membrane disruption. LDH levels were significantly lower for *Δngo-lig E*^*kan*^ on the rHVE cells relative to infection with the *wt* strain (Figure 5), mirroring the trend observed when the invasion depth was calculated from the CLSM images (Figure 4 (d)). Interestingly, although the amount of LDH released by cells infected with the overexpressor *opaB-ngo-lig E-his*^*kan*^ was also significantly lower than that of *wt*, this decrease was not as large as that caused by the *Δngo-Lig E*^*kan*^ mutant.

**Figure 5.**
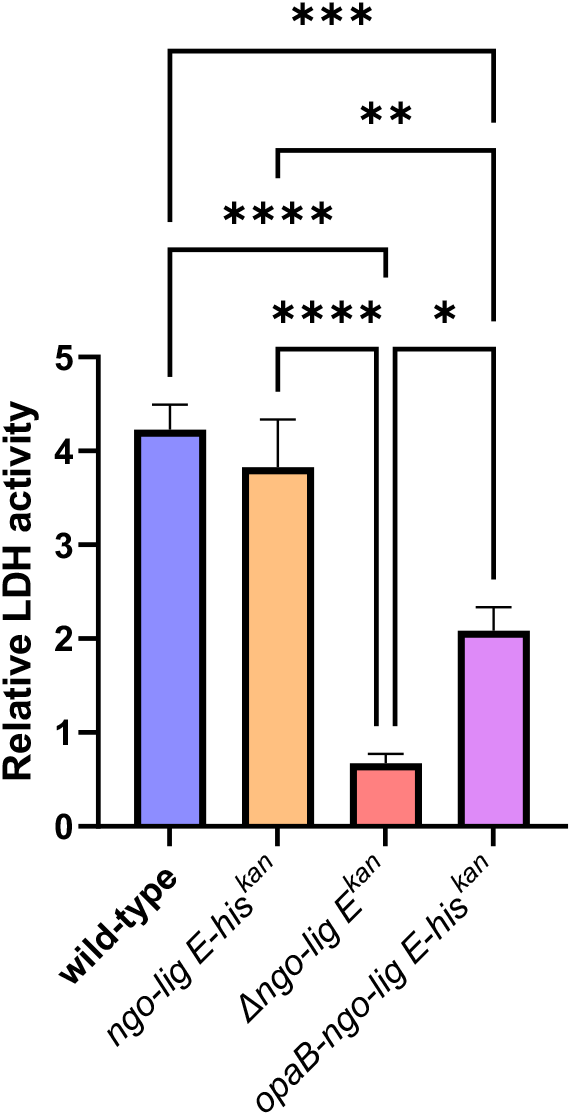
Quantification of lactase dehydrogenase (LDH) release in the supernatant after N. gonorrhoeae infection of rHVE tissue. Relative LDH activity was obtained via absorbance at 490 nm relative to the absorbance of blank media and corrected for by the background absorbance at 680 nm. Points are the mean of values from three technical replicates of each biological replicate and error bars represent the standard error of the mean. Significance values are given as * p ≤ 0.05; ** p ≤ 0.01; *** p ≤ 0.001; **** p ≤ 0.0001 Comparisons which showed no significant differences (p > 0.05) are not indicated.

### Ngo-Lig E is important for microcolony formation in *Neisseria gonorrhoeae*

To determine if the observed differences in biofilm morphology were attributable to altered microcolony formation, SEM microscopy on the *N. gonorrhoeae* strains during exponential-phase growth was performed. These images showed that *Δngo-lig E*^*kan*^ forms markedly fewer and smaller microcolonies compared to *wt N. gonorrhoeae* (Figure 6(a) and Figure S4), and that these covered significantly lower surface areas (Figure 6(b)). Furthermore, the microcolonies formed by *ngo-lig E*^*kan*^ were slightly more dispersed and ‘loose’ compared to the cohesive microcolonies formed by the *wt* strain. Meanwhile the his-tagged control (*ngo-lig E-his*^*kan*^) microcolonies covered a similar total surface area to the *wt* (Figure 6(b)). To evaluate the impact of increased exDNA on microcolony formation, microcolonies formed by the *nuc* deletion strain (*Δnuc*^*kan*^) were imaged. Deletion of *nuc* has been previously demonstrated to increase biofilm formation in *N. gonorrhoeae* (Steichen *et al*., 2011), and here, we showed that the microcolonies formed by this mutant were similar in size to that formed by *wt*, albeit slightly denser and more compact (Figure 6(a)). Furthermore, the *Δnuc*^*kan*^ cells also adhered to one another more closely in what appears to be a thicker extracellular matrix (ECM).

**Figure 6.**
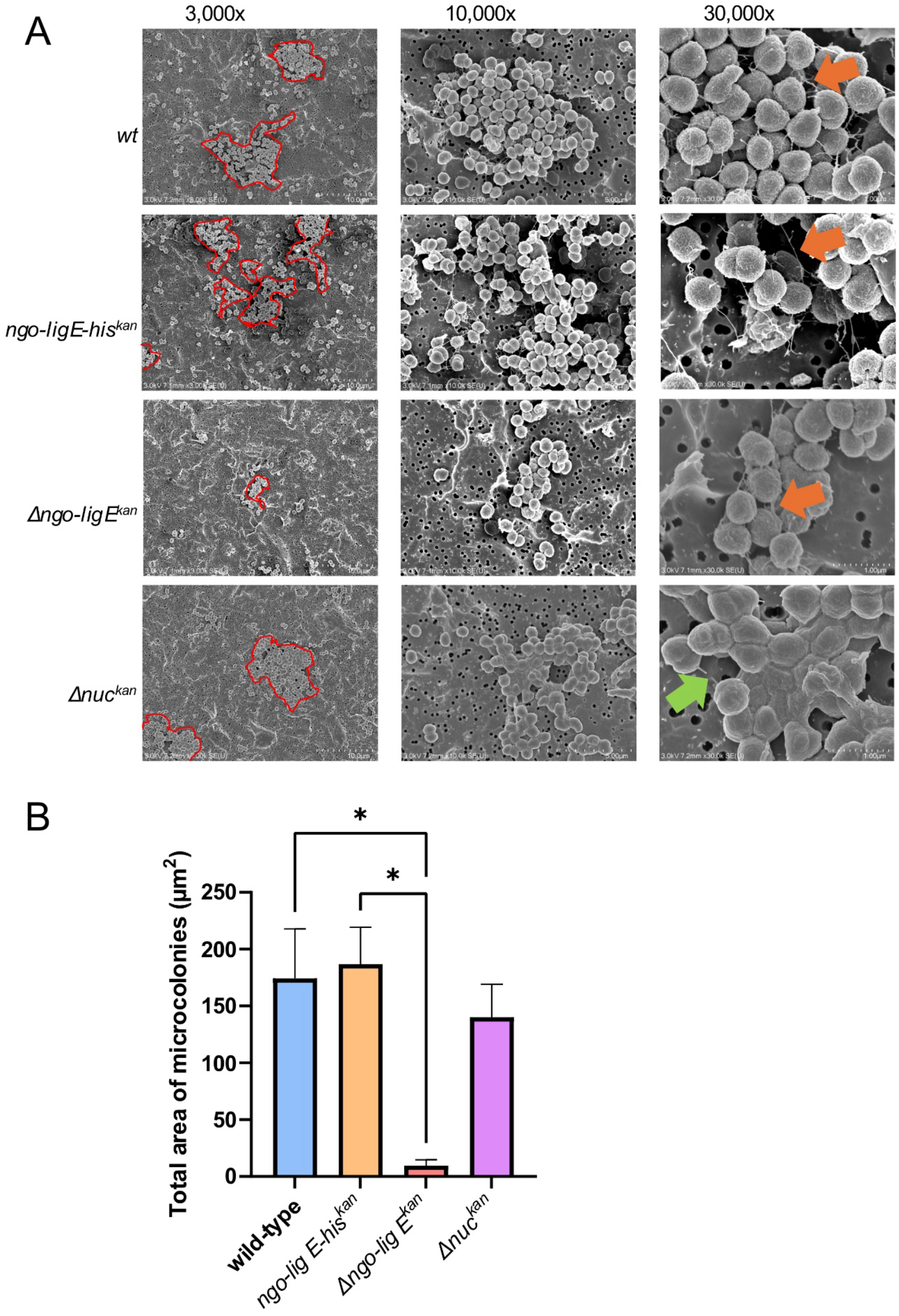
N. gonorrhoeae microcolony formation (a) Representative SEM images of N. gonorrhoeae microcolonies formed on 0.2 µm pore size filter paper during the exponential phase of growth (9 hours). The approximate outline of microcolonies in the field-of-view are shown in red and were annotated manually after image collection. Orange arrows indicate pili filaments, while the green arrow points at a similar extended filamentous structure. Additional supporting fields-of-view can be found in Fig S4. (b) Quantification of the average total area covered by the microcolonies in each field-of-view via ImageJ (Schneider et al., 2012). Points are the mean of values of the total surface area covered in each field-of view and error bars represent the standard error of the mean. Significance values are given as * p ≤ 0.05. Comparisons which showed no significant differences (p > 0.05) are not indicated.

Images at a higher magnification (x30,000) showed piliation filaments that seemed to tether individual bacteria to each other in the microcolonies for both *wt* and the his-tagged control, *ngo-lig E-his*^*kan*^ (Figure 6 (a), indicated via orange arrows). Although also present in *Δngo-lig E*^*kan*^, these piliation filaments were not as extensive when *ngo-lig E* was disrupted. Interestingly, the extensions observed projecting between cells in the *Δnuc*^*kan*^ mutant were thicker than the pili filaments observed in the other variants (Figure 6 (a), indicated by the green arrow), which could be due to a covering of ECM.

## Discussion

In a previous study, we showed via an end-point crystal violet assay that the disruption of the DNA ligase Ngo-Lig E negatively affected biofilm formation by *N. gonorrhoeae* (Pan *et al*., 2024). Here, we have expanded on this finding by culturing the same mutants under constant shear forces and in continuous medium flow in CDC Biofilm Reactors®. The morphology of the resulting biofilms was assessed and demonstrated a marked reduction in colonisation and biofilm formation on artificial surfaces when *ngo-lig E* was disrupted compared to the *wt* and the *ngo-lig E-his*^*kan*^ mutant. The latter strain served as a control for the insertion of the kanamycin resistance marker which was used to construct the deletion and overexpressing strains, and indicated that the resistance cassette was not responsible for the observed phenotype. Conversely, overexpression of *ngo-lig E* (*opa-lig E-his*^*kan*^) formed denser and thicker biofilms which were more compact relative to *wt*. We previously demonstrated a 90-fold upregulation of *ngo-lig E* expression in this strain, indicating that Ngo-Lig E activity contributed positively to biofilm formation (Pan *et al*., 2024)). Our previous work (Pan *et al*., 2024) also revealed no differences in growth rates between any of the *ngo-lig E* variants and *wt*, and since the inocula were normalised before each experiment, the observed differences in biofilm formation were not due to differences in initial cell density or growth rates.

Consistent with the phenotype on artificial surfaces, *ngo-lig E* deletion also causes defects in *N. gonorrhoeae* attachment to human cells and invasion into tissue. Previous results from our group showed decreased adhesion of the *Δngo-lig E*^*kan*^ mutant on a monolayer of ME-180 human cervical cells (Pan *et al*., 2024). Here, we also show the reduced capabilities of the same mutant to invade and migrate into reconstituted 3-D HVE tissue, which in turn equates to less cellular damage by the deletion mutant relative to *wt N. gonorrhoeae*. Instead, *Δngo-lig E*^*kan*^ accumulated on the upper surface of the tissue, which appeared healthier and less perforated than in the *wt*-infected sections. This has important implications on the extent of *N. gonorrhoeae* infection in the human host, as *N. gonorrhoeae* invasion and transcytosis into tissues may lead to disseminated infections after crossing the subepithelial space (Edwards & Apicella, 2004). Although the overexpressor *opa-ngo-lig E-his*^*kan*^ appeared to follow this trend as well, we note that most of the bacteria were closer to the membrane, and we speculate that this could be indicative of deeper invasion past the membrane into the supernatant. We predict that due to the decreased ability of *Δngo-lig E*^*kan*^ to attach to the surfaces of cells, the strain is less able to invade human cells and cause damage. However, as we were only able to obtain end-point images after 16-17 h of infection, we did not capture the step-by-step process of internalisation and transcytosis of the other mutants into the host cells as it occurred (*e*.*g*. via time-lapse microscopy).

Our observations of the consequences of *ngo-lig E* deletion on biofilm morphology are consistent with our hypothesis that this ligase acts on exDNA to form high molecular weight substrates that better contribute to initial biofilm formation by overcoming any repulsive forces between the ECM and the surface (Maier, 2021). Unlike many other bacterial biofilms, exDNA is likely the primary structural biopolymer of gonococcal biofilms, with dsDNA being a critical component of mature *N. gonorrhoeae* biofilms (Zweig *et al*., 2014; Kouzel *et al*., 2015). Work by Bender *et al*. has demonstrated that for *N. gonorrhoeae* larger pieces of DNA (>3000 bp) are more likely to have a high concentration gradient outside the colony and integrate into the biofilm (Bender *et al*., 2022). We hypothesise that extracellular Ngo-Lig E is important for repairing breaks in free exDNA fragments, increasing their length and integrity and allowing them to be retained outside the colonies and therefore contribute to the pool of exDNA that stabilises the extracellular biofilm matrix of *N. gonorrhoeae*. This action would counteract the activity of the extracellular Nuc, which remodels gonococcal biofilms through its cleavage and degradation of exDNA (Steichen *et al*., 2011) and suggests that some level of interplay or regulatory control between these two opposing activities is likely. Here, we show that the *nuc* deletion strain formed microcolonies of similar size to *wt N. gonorrhoeae*; however, these microcolonies appeared more globular, with filamentous extensions that seemed to indicate thicker ECM. It is possible that this observed morphology was due to the inability of the *nuc* deletion to regulate exDNA content, and suggests that in the *wt* strain, Nuc works in conjunction with Ngo-Lig E to manipulate exDNA, and thus optimise and modulate the architecture of the biofilm. We also question if the increase in exDNA length and integrity performed by Ngo-Lig E allows the exDNA to interact with other DNA binding proteins, which may assemble to create a more stable framework for the biofilm to build on.

Here, we have shown that Ngo-Lig E affects biofilm formation, potentially via its activity on exDNA, which in turn affects the adhesion of *N. gonorrhoeae* to human cells, and their subsequent invasion and damage. Based on these observations, we consider extracellular Ngo-Lig E to be important for the pathogenicity and virulence of *N. gonorrhoeae*, making it an appealing target for future drug design against this incredibly resistant bacterium. Despite this, many questions about Ngo-Lig E remain, including its specific cellular location, its regulation and its direct consequences for the exDNA fraction. The presence of an N-terminal signal peptide indicates that Ngo-Lig E is transported to the periplasmic space, however, it may be further transported to the extracellular milieu via membranous blebs that contribute to the ECM of *N. gonorrhoeae* biofilms (Dorward & Garon, 1989; Dorward *et al*., 1989; Steichen *et al*., 2008). Additionally, it remains unknown if Ngo-Lig E is required for biofilm maintenance, or if it is more important during early biofilm formation when exDNA is most critical (Zweig *et al*., 2014). Answering this question would require quantifying the amount and arrangement of exDNA in the ECM when *ngo-lig E* is disrupted (i.e via exDNA staining); this was attempted in the present study, however we encountered difficulties in its visualisation after growth in the Biofilm Reactors, potentially due to the constant stirring that may damage the exDNA.

A final area of interest is how Ngo-Lig E activity, and especially a potential influence on DNA size may affect gene transfer, which is more frequent in early biofilms (Kouzel *et al*., 2015). This would involve studying biofilm formation by the different gonococcal mutants at different timepoints.

### New tools and methods for understanding *Neisseria gonorrhoeae* biofilm and pathogenicity

Previously, our group had used a crystal violet assay to demonstrate the importance of Ngo-Lig E on biofilm formation, which although rapid and convenient, was an indirect method that assessed static biofilms (Pan *et al*., 2024). Other more advanced techniques that have been used to study *N. gonorrhoeae* biofilms include the growth of the bacteria on glass or patterned silicone coverslips (the latter to study the effect of surface topography) in continuous flow chambers or cells, or even static growth on glass dishes (Greiner *et al*., 2005; Falsetta *et al*., 2009; Falsetta *et al*., 2011; Steichen *et al*., 2011; Kwiatek *et al*., 2014; Kouzel *et al*., 2015; Oldewurtel *et al*., 2015; Płaczkiewicz *et al*., 2019). While such methods have greatly increased our understanding of gonococcal biofilm formation, they often require bespoke laboratory equipment, and can be subject to technical complications such as bubble formation in the flow channels that disrupt cellular adhesion

Here, we report the first use of a commercially-available CDC Biofilm Reactor® to study *N. gonorrhoeae* biofilms, involving the continuous flow of fresh medium controlled via a periplasmic pump into a growth chamber filled with retrievable coupons that the bacteria can adhere to, while maintaining a constant shear force across the surface (Kocot *et al*., 2021). Adhesion of *N. gonorrhoeae* to polycarbonate coupons was not as extensive as we had anticipated, however we attribute this to the polycarbonate material used. This was readily available in our laboratory, but has not been widely used for *N. gonorrhoeae* biofilm studies, with glass being the preferred substrate for *N. gonorrhoeae* surface adhesion for other reports (Greiner *et al*., 2005; Falsetta *et al*., 2009; Falsetta *et al*., 2011; Steichen *et al*., 2011; Kwiatek *et al*., 2014; Płaczkiewicz *et al*., 2019). In addition, the constant stirring of medium may have damaged exDNA, which since it is a major constituent of *N. gonorrhoeae* biofilms, would also affect the adhesion and biofilm formation in this setting. Despite this, we believe that with further optimisation, CDC Biofilm Reactors® offer great promise for further studies of biofilm formation in *N. gonorrhoeae*, especially if a glass surface and slower shear forces are used.

We also showed the importance of a 3-D model for studying bacterial-host interactions, which allowed us to examine the effects of Ngo-Lig on *N. gonorrhoeae* migration into host tissues; a phenotype not observable in a cell monolayer. Common techniques employed by other groups involve the use of a primary cell line or biopsy samples which can be directly coated onto glass coverslips for easy microscopy (Greiner *et al*., 2005; Płaczkiewicz *et al*., 2019). The search for appropriate 3-D models for *N. gonorrhoeae* study is of increasing interest, with one particular group developing a model for this purpose using porcine small intestinal submucosa as a scajold (Heydarian *et al*., 2019). Until models like this are readily available however, the SkinEthic HVE™ tissue model used here from Episkin (Lyon, France) provides a good substitute as it is easy to obtain, reproducible and provides a more biologically-relevant model with different tissue cell types and morphologies.

## Conclusion

The DNA ligase, Lig E, is present in many bacteria like *N. gonorrhoeae* that form exDNA-dependent biofilms. Here we show that Lig E from *N. gonorrhoeae* (Ngo-Lig E) influences the formation of gonococcal biofilms and microcolonies on artificial surfaces, as well as the invasion into and damage of 3-D reconstituted HVE tissue. We propose that Ngo-Lig E may be acting on fragmented exDNA in the extracellular space of *N. gonorrhoeae*, which is conducive for microcolony formation and proto-biofilm interactions to occur. Future directions include studying the role of Ngo-Lig E at different stages of gonococcal biofilm formation, as well as investigation into the potential interplay between Ngo-Lig E and Nuc on exDNA-mediated biofilm remodelling. Regardless, the results presented in this report highlight the importance of Ngo-Lig E on the virulence and pathogenicity of *N. gonorrhoeae*. We predict that this may open up new avenues and pathways for targeting not only *N. gonorrhoeae*, but other human pathogens that express this minimal ligase, potentially finding a way to target extensive biofilm formation in many clinical settings.

## Acknowledgements

We would like to thank Fiona Clow and Fiona Radcliff from the Radcliff Laboratory (University of Auckland) for providing us with the pEG2 plasmid, as well as Adam Jones (School of Dentistry, Cardiff University) for guidance with histological processing of the tissue. We would also like to extend our thanks to Helen Turner (University of Waikato) for performing the critical point drying process for SEM microscopy and Marc Isaacs (Bioimaging Hub, Cardiff University) for help with wax microtomy and slide preparation. This work was supported by the Maurice Wilkins Centre Flexible Research Programme (Category 4) and the Cardiff University and the University of Waikato Strategic International Partnership Collaborative Seed Fund. AW is supported by a Rutherford Discovery Fellowship (20-UOW-004). JP is supported by a University of Waikato Doctoral Scholarship.

## Author contributions

JP led the experimental work including the construction, growth, biofilm formation in CDC Biofilm Reactors^®^, rHVE infection and confocal visualisation of the *N. gonorrhoeae* mutants. AA provided expertise and guidance and assisted with setting up the CDC Biofilm Reactors^®^, rHVE infections, as well as confocal imaging of the coupons and tissue slides. The study and experimental design of the work was conceptualised by JP, AW, DW and JH.

## References

Bender, N., Hennes, M., & Maier, B. (2022). Mobility of extracellular DNA within gonococcal colonies. Biofilm, 4, 100078.

Christodoulides, M., Everson, J. S., Liu, B. L., Lambden, P. R., Watt, P. J., Thomas, E. J., & Heckels, J. E. (2000). Interaction of primary human endometrial cells with Neisseria gonorrhoeae expressing green fluorescent protein. Molecular Microbiology, 35(1), 32–43.

Dillard, J. P. (2011). Genetic manipulation of Neisseria gonorrhoeae. Current Protocols in Microbiology, Chapter 4, Unit4A.2-Unit4A.2.

Dorward, D. W., & Garon, C. F. (1989). DNA-binding proteins in cells and membrane blebs of Neisseria gonorrhoeae. Journal of Bacteriology, 171(8), 4196–201.

Dorward, D. W., Garon, C. F., & Judd, R. C. (1989). Export and intercellular transfer of DNA via membrane blebs of Neisseria gonorrhoeae. Journal of Bacteriology, 171(5), 2499–505.

Edwards, J. L., & Apicella, M. A. (2004). The molecular mechanisms used by Neisseria gonorrhoeae to initiate infection differ between men and women. Clinical Microbiology Reviews, 17(4), 965–81.

Elmros, T., Burman, L. G., & Bloom, G. D. (1976). Autolysis of Neisseria gonorrhoeae. Journal of Bacteriology, 126(2), 969–76.

Falsetta, M. L., Bair, T. B., Ku, S. C., Vanden Hoven, R. N., Steichen, C. T., McEwan, A. G., Jennings, M. P., & Apicella, M. A. (2009). Transcriptional profiling identifies the metabolic phenotype of gonococcal biofilms. Infection and Immunity, 77(9), 3522–32.

Falsetta, M. L., Steichen, C. T., McEwan, A. G., Cho, C., Ketterer, M., Shao, J., Hunt, J., Jennings, M. P., & Apicella, M. A. (2011). The composition and metabolic phenotype of Neisseria gonorrhoeae biofilms. Frontiers in Microbiology, 2, 75–75.

Greiner, L. L., Edwards, J. L., Shao, J., Rabinak, C., Entz, D., & Apicella, M. A. (2005). Biofilm formation by Neisseria gonorrhoeae. Infection and Immunity, 73(4), 1964–1970.

Hebeler, B. H., & Young, F. E. (1975). Autolysis of Neisseria gonorrhoeae. Journal of Bacteriology, 122(2), 385–92.

Heydarian, M., Yang, T., Schweinlin, M., Steinke, M., Walles, H., Rudel, T., & Kozjak-Pavlovic, V. (2019). Biomimetic human tissue model for long-term study of Neisseria gonorrhoeae infection. Frontiers in Microbiology, 10.

Heydorn, A., Nielsen, A. T., Hentzer, M., Sternberg, C., Givskov, M., Ersbøll, B. K., & Molin, S. (2000). Quantification of biofilm structures by the novel computer program COMSTAT. Microbiology (Reading), 146 (Pt 10), 2395–2407.

Juneau, R. A., Stevens, J. S., Apicella, M. A., & Criss, A. K. (2015). A thermonuclease of Neisseria gonorrhoeae enhances bacterial escape from killing by neutrophil extracellular traps. The Journal of Infectious Diseases, 212(2), 316–324.

Kocot, A., Wróblewska, B., & Cabo, M. (2021). Operational culture conditions determinate benzalkonium chloride resistance in L. monocytogenes-E. coli dual species biofilms. International Journal of Food Microbiology, 360, 109441.

Kouzel, N., Oldewurtel, E. R., & Maier, B. (2015). Gene transfer ejiciency in gonococcal biofilms: Role of biofilm age, architecture, and pilin antigenic variation. Journal of Bacteriology, 197(14), 2422–2431.

Kwiatek, A., Bacal, P., Wasiluk, A., Trybunko, A., & Adamczyk-Poplawska, M. (2014). The dam replacing gene product enhances Neisseria gonorrhoeae FA1090 viability and biofilm formation. Frontiers in Microbiology, 5(712).

Magnet, S., & Blanchard, J. S. (2004). Mechanistic and kinetic study of the ATP-dependent DNA ligase of Neisseria meningitidis. Biochemistry, 43(3), 710–717.

Maier, B. (2021). How physical interactions shape bacterial biofilms. Annual Review of Biophysics, 50(1), 401–417.

Malfa, P., Brambilla, L., Giardina, S., Masciarelli, M., Squarzanti, D. F., Carlomagno, F., & Meloni, M. (2023). Evaluation of antimicrobial, antiadhesive and co-aggregation activity of a multi-strain probiotic composition against different urogenital pathogens. International Journal of Molecular Sciences, 24(2), 1323.

McCormack, W. M. (1981). Clinical spectrum of infection with Neisseria gonorrhoeae. Sexually Transmitted Diseases, 8(4), 305–307.

Morse, S. A., & Bartenstein, L. (1974). Factors ajecting autolysis of Neisseria gonorrhoeae. Experimental Biology and Medicine (Maywood, N.J.), 145(4), 1418–1421.

Murga, R., Stewart, P. S., & Daly, D. (1995). Quantitative analysis of biofilm thickness variability. Biotechnology and Bioengineering, 45(6), 503–510.

Nassar, R., Nassar, M., Senok, A., & Williams, D. (2023). Phytic acid demonstrates rapid antibiofilm activity and inhibits biofilm formation when used as a surface conditioning agent. Microbiology Spectrum, 11(3), e00267–23.

Nowicki, B. (1985). Multiparameter representation of surface roughness. Wear, 102(3), 161–176.

Oldewurtel, E. R., Kouzel, N., Dewenter, L., Henseler, K., & Maier, B. (2015). Differential interaction forces govern bacterial sorting in early biofilms. eLife, 4, e10811.

Pan, J., Lian, K., Sarre, A., Leiros, H.-K. S., & Williamson, A. (2021). Bacteriophage origin of some minimal ATP-dependent DNA ligases: a new structure from Burkholderia pseudomallei with striking similarity to Chlorella virus ligase. Scientific Reports, 11(1), 18693.

Pan, J., Singh, A., Hanning, K., Hicks, J., & Williamson, A. (2024). A role for the ATP-dependent DNA ligase lig E of Neisseria gonorrhoeae in biofilm formation. BMC Microbiology, 24(1), 29.

Płaczkiewicz, J., Adamczyk-Popławska, M., Lasek, R., Bącal, P., & Kwiatek, A. (2019). Inactivation of genes encoding MutL and MutS proteins influences adhesion and biofilm formation by Neisseria gonorrhoeae. Microorganisms, 7(12), 647.

Schneider, C. A., Rasband, W. S., & Eliceiri, K. W. (2012). NIH Image to ImageJ: 25 years of image analysis. Nature Methods, 9(7), 671–675.

Steichen, C. T., Cho, C., Shao, J. Q., & Apicella, M. A. (2011). The Neisseria gonorrhoeae biofilm matrix contains DNA, and an endogenous nuclease controls its incorporation. Infection and Immunity, 79(4), 1504–1511.

Steichen, C. T., Shao, J. Q., Ketterer, M. R., & Apicella, M. A. (2008). Gonococcal cervicitis: a role for biofilm in pathogenesis. The Journal of Infectious Diseases, 198(12), 1856–1861.

Tapsall, J. W. (2005). Antibiotic resistance in Neisseria gonorrhoeae. Clinical Infectious Diseases, 41(Supplement_4), S263–S268.

Unemo, M., & Shafer, W. M. (2014). Antimicrobial resistance in Neisseria gonorrhoeae in the 21st century: Past, evolution, and future. Clinical Microbiology Reviews, 27(3), 587–613.

Vorregaard, M. (2008). Comstat2 - a modern 3D image analysis environment for biofilms. In Informatics and Mathematical Modelling. Kongens Lyngby, Denmark: Technical University of Denmark.

Williamson, A., Hjerde, E., & Kahlke, T. (2016). Analysis of the distribution and evolution of the ATP-dependent DNA ligases of bacteria delineates a distinct phylogenetic group ‘Lig E’. Molecular Microbiology, 99(2), 274–90.

Williamson, A., Rothweiler, U., & Leiros, H. K. (2014). Enzyme-adenylate structure of a bacterial ATP-dependent DNA ligase with a minimized DNA-binding surface. Acta Crystallographica Section D: Biological Crystallography, 70(Pt 11), 3043–56.

World Health Organization. (2024). Global sexually transmitted infections programme: Diagnostics for gonococcal antimicrobial resistance. from https://www.who.int/teams/global-hiv-hepatitis-and-stis-programmes/stis/testing-diagnostics/diagnostics-for-gonococcal-antimicrobial-resistance.

Zola, T. A., Strange, H. R., Dominguez, N. M., Dillard, J. P., & Cornelissen, C. N. (2010). Type IV secretion machinery promotes ton-independent intracellular survival of Neisseria gonorrhoeae within cervical epithelial cells. Infection and Immunity, 78(6), 2429–37.

Zweig, M., Schork, S., Koerdt, A., Siewering, K., Sternberg, C., Thormann, K., Albers, S. V., Molin, S., & van der Does, C. (2014). Secreted single-stranded DNA is involved in the initial phase of biofilm formation by Neisseria gonorrhoeae. Environmental Microbiology, 16(4), 1040–52.

