## Supplementary for "Influence of the ATP-dependent DNA ligase, Lig E, on *Neisseria gonorrhoeae* microcolony and biofilm formation"

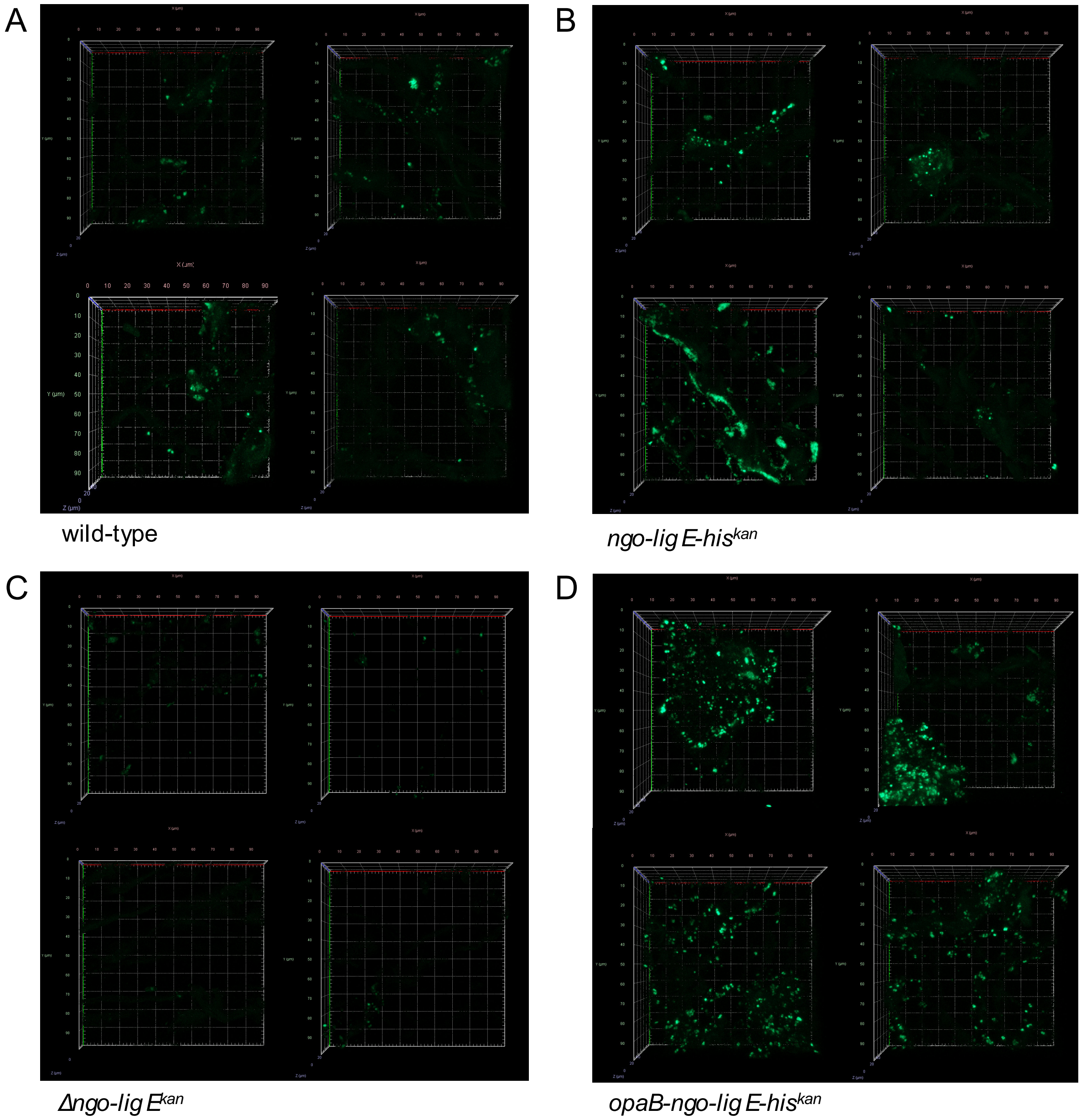


Figure S1. Additional CLSM z-stack images of representative fields-of-view of *N. gonorrhoeae* pEG2 (expressing sfGFP) biofilm formation and adhesion on polycarbonate coupons in CDC Biofilm Reactors® (x40 magnification, em:480 nm, exc:505 nm).

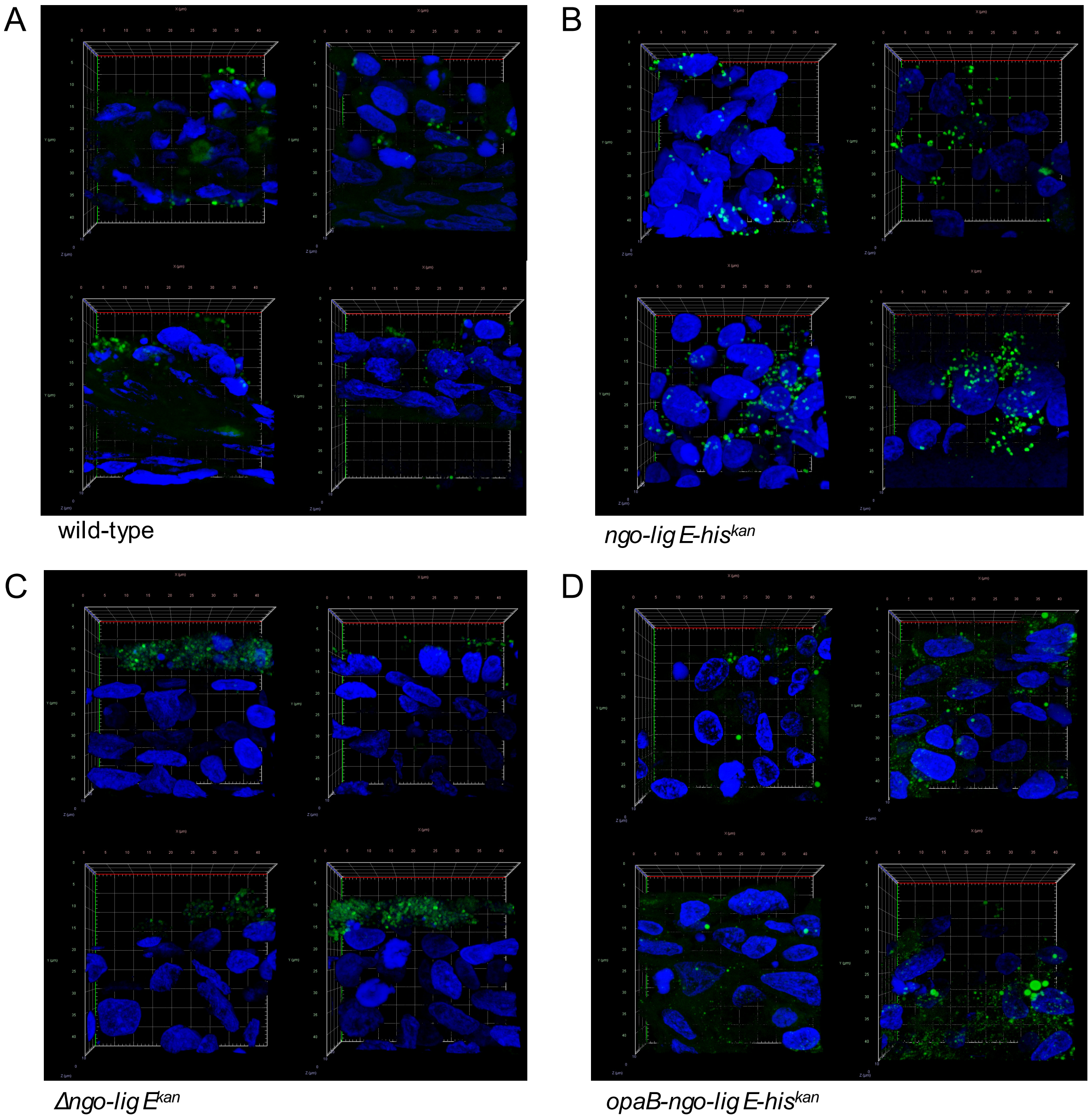


Figure S2. Additional CLSM z-stack images (x63 objective magnification, zoom setting 3) of representative fields-of-view of the infection and invasion of SkinEthic™ HVE cells by *N. gonorrhoeae*. *N. gonorrhoeae* pEG2 (expressing sfGFP) is shown in green (exc: 488 nm, em: 519 nm) and the nuclei of the rHVE cells in blue (exc: 405 nm, em: 449 nm).

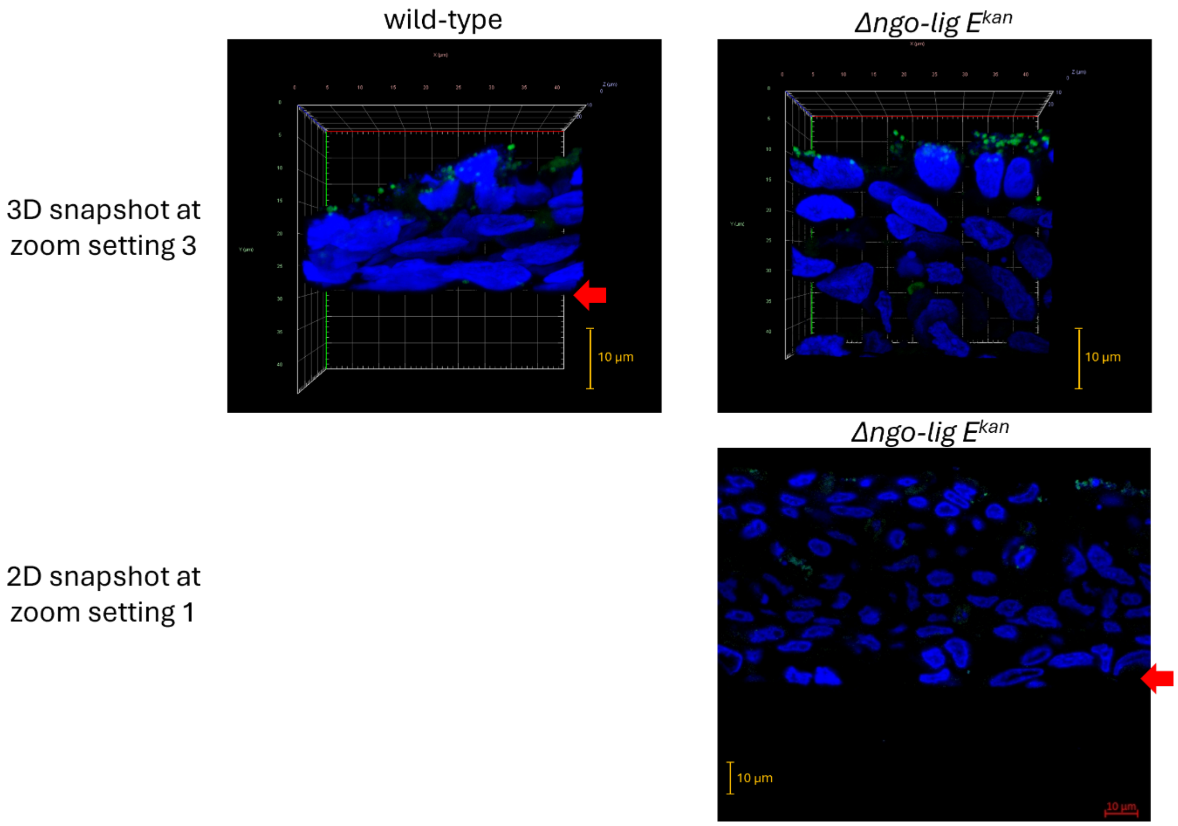


Fig S3. CLSM images (x63 objective magnification) of SkinEthic™ HVE cells (blue, exc: 405 nm, em: 449 nm) infected with *N. gonorrhoeae* pEG2 (green, exc: 488 nm, em: 519 nm) showing the extent of tissue damage. The top row shows the 3-D snapshot of *wt* and *Δngo-lig* $E^{kan}$ *N. gonorrhoeae* at zoom setting 3 (final zoom setting used), showing the depth and length of the remaining intact tissue, with the tissue membrane visible for *wt* at that setting (red arrow). The bottom row shows a 2-D snapshot of the same *Δngo-Lig* $E^{kan}$ infected tissue slide at zoom setting 1 which finally shows the membrane of the tissue (red arrow).

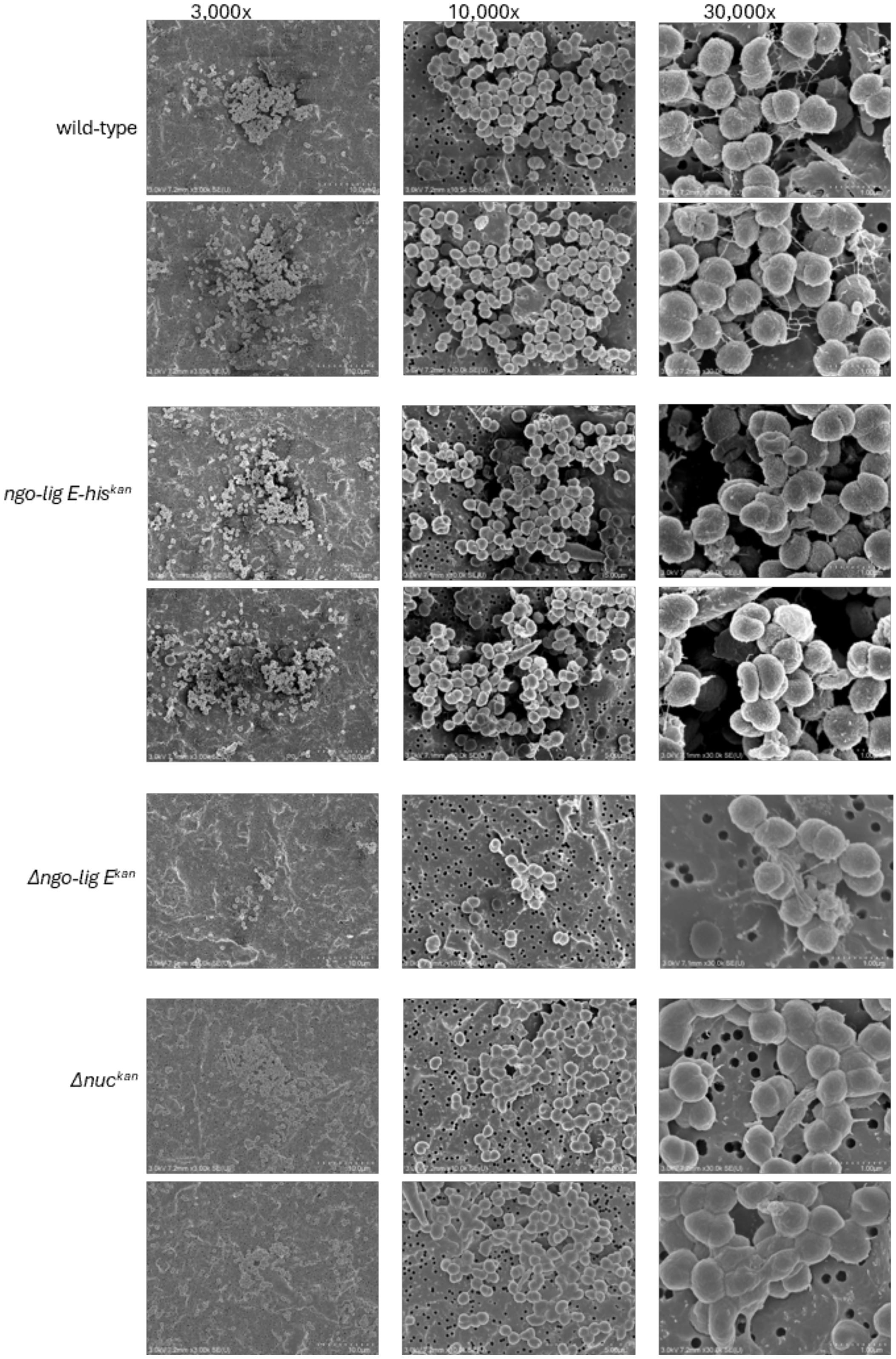


Figure S4. Additional supporting SEM images of *N. gonorrhoeae* microcolonies formed on 0.2 µm pore size filter paper during the exponential phase of growth (9 hours). Note, only one other field-of-view is shown for *Δngo-lig E*$^{kan}$ due to the lack of microcolonies formed by this mutant.
